# Phenotype Heritability in holobionts: An Evolutionary Model

**DOI:** 10.1101/2020.05.05.079137

**Authors:** Saúl Huitzil, Santiago Sandoval-Motta, Alejandro Frank, Maximino Aldana

## Abstract

Many complex diseases are expressed with high incidence only in certain populations. Genealogy studies determine that these diseases are inherited with a high probability. However, genetic studies have been unable to identify the genomic signatures responsible for such heritability, as identifying the genetic variants that make a population prone to a given disease is not enough to explain its high occurrence within the population. This gap is known as the missing heritability problem. We know that the microbiota plays a very important role in determining many important phenotypic characteristics of its host, in particular, the complex diseases for which the missing heritability occurs. Therefore, when computing the heritability of a phenotype it is important to consider not only the genetic variation in the host but also in its microbiota. Here we test this hypothesis by studying an evolutionary model based on gene regulatory networks. Our results show that the holobiont (the host plus its microbiota) is capable of generating a much larger variability than the host alone, greatly reducing the missing heritability of the phenotype. This result strongly suggests that a considerably large part of the missing heritability can be attributed to the microbiome.

## 1 Introduction

One of the main objectives of the Human Genome Project, the HapMap Project, and later of the GWAS studies, has been to understand the genetic architecture Responsible for the emergence of complex diseases [6,7,34]. For a long time, through genealogy studies, it has been estimated that such complex diseases have a strong heritability determined by genetic components [23]. However, this type of study has the inconvenience that it is limited to a small number of individuals. By contrast, thanks to high-throughput techniques such as GWAS, the size of the population under study could be considerably increased because the degree of kinship between individuals can be determined through their genetic variants (SNPs).

The first GWAS study was performed in 2002 [25]. Since then, this method has been very successful in associating thousands of genetic variants with different phenotypic traits [4]. However, the effect of these genetic variants on a particular phenotype is very small [22] and, consequently, they can explain only a small fraction of the heritability of the associated phenotype. This is because the fraction of phenotypic variance that can be attributed to additive genetic factors is considerably smaller than expected when we consider the genetic variants associated with a given phenotype by GWAS [13].

The gap between the heritability observed through genealogy studies and the one measured from GWAS is known as the *missing heritabilit*y, and for some phenotypes can be as large as 60% [33]. A typical example is human height. On the one hand, familial studies have estimated that this phenotype has a heritability larger than 80%. On the other hand, through GWAS about 700 SNPs have been associated with human height. However, these SNPs can explain only 20% of its heritability [37]. The most used strategy to fill up the missing heritability gap has been to find more and more genetic variants associated with a particular phenotype in order to increase its estimated heritability [13]. In 2010, Yang et al. computed the heritability produced by 29,4831 SNPs, which allowed them to estimate the heritability of human height around 45% [38]. Nonetheless, if the missing heritability problem were to be solved by increasing the number of SNPs, then we would have to consider so many genetic variants that we would still be unable to understand the genetic architecture behind the emergence of complex diseases and phenotypes. Several solutions have been proposed, such as taking into account epigenetic and epistatic effects, the effect of non-coding RNAs in gene regulation, or an overestimation of the heritability itself [22, 23, 29, 41, 14, 10]. However, there is no consensus yet in the scientific community as to how to solve this problem.

During the last decade, numerous studies have shown the importance of the microbiome in determining important phenotypic traits of the host organism [3]. The microbiota composition and functionality have been correlated with many complex diseases, allergies, neurological disorders, metabolism of antibiotics and other drugs, to mention just a few [5, 17, 30, 19, 36]. Additionally, there is evidence suggesting that phenotypic traits can be transmitted (or inherited) from the parents to their offspring through the microbiota [18]. For instance, the phylogenetic congruence (phylosymbiosis) between the host organism and its microbiota suggests that a set of microorganisms have been inherited throughout evolution [21, 24]. Furthermore, it is known that the metabolic response of several host organisms to the microbiota is conserved throughout evolution across different vertebrates, including the zebrafish, mice, and humans [26]. The microbiota participates in practically all important biological tasks of the host [5]. Given the strong symbiotic interactions between the host and its microbiota, and the evidence showing that microorganisms can be transmitted both vertically and horizontally from parents to offspring, Rosenberg, and his collaborators formulated the hypothesis of the holobiont as a unit of selection in evolution [39]. This hypothesis states that in order to understand the evolution of the phenotypic traits of a given organism one has to consider the hologenome (the genome of the host and the genomes of all of its microbes) as a unit of selection [39]. This means that changes in the phenotype of a host organism can be produced by mutations in the host’s genome, in the genes of its microbes (the host’s microbiome), or in both, and to understand these phenotypic changes one has to consider genetic variation in the entire hologenome. This may be particularly true for complex phenotypes for which missing heritability exists. Therefore, it is reasonable to assume that, when computing the heritability of a phenotype, the genetic variants of the entire hologenome have to be taken into account. This hypothesis was proposed in previous works [28, 32], but to our knowledge, no measurements have been performed so far validating or refuting this idea. Here we present an evolutionary model, based on Boolean networks, showing quantitatively that a large part of the missing heritability is recovered when the genetic variants are computed using the entire hologenome rather than just the genome of the host.

## 2 Boolean Network model

We choose to work with Boolean networks as representations of the genomes of both the host and its microbes. These networks were proposed by S. Kauffman in 1969 as models for gene regulation, and have proven to reproduce the gene expression patterns observed experimentally for several organisms [2,9]. A Boolean network consists of *N* nodes representing the genes in the genome. The state of expression of these genes is encoded in a set of variables {*g*_1_, *g*_2_, … *g*_*N*_} such that *g*_*n*_ = 1 if the *n*^*th*^ gene is expressed, or *g*_*n*_ = 0 if it is not expressed. The value of each *g*_*n*_ is determined by a particular set of *k*_*n*_ other genes in the network, which we represent as 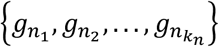 (see Fig.1a). We will refer to this set as the *inputs* or *regulators* of *g*_*n*_, and to *k*_*n*_ as its *input connectivity* which can be different from one gene to another. Once each gene in the networks has been assigned with a set of regulators, the network dynamics is determined by the simultaneous update of the state of all the genes according to the equation

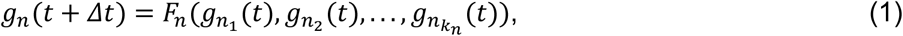

where 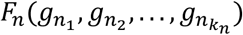 is a Boolean function of *k*_*n*_ variables and *Δt* is the average time it takes for a gene to respond to changes in its regulators. For networks of real organisms, the set of regulators of each gene *g*_*n*_ is determined from experimental observations and its Boolean function is constructed by hand according to the activating or inhibitory nature of its regulators (see Fig. 1b). Here we are interested in the general evolutionary properties of holobionts and not on any particular organism. Therefore, we are going to work with Boolean networks constructed in the following way: (a) the number *k*_*n*_ of inputs of each gene *g*_*n*_will be randomly chosen from a predefined probability function *P*_*I*_(*k*), which has average *K* and variance 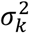; (b) the *k*_*n*_ inputs of each gene are chosen randomly from anywhere in the in the network with uniform probability; (c) each Boolean function *F*_*n*_ will be constructed randomly such that for each of the 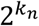 configurations of its *k*_*n*_ arguments, *F*_*n*_ = 1 with probability *p* and *F*_*n*_ = 0 with probability 1 − *p*.

**Fig. 1.**
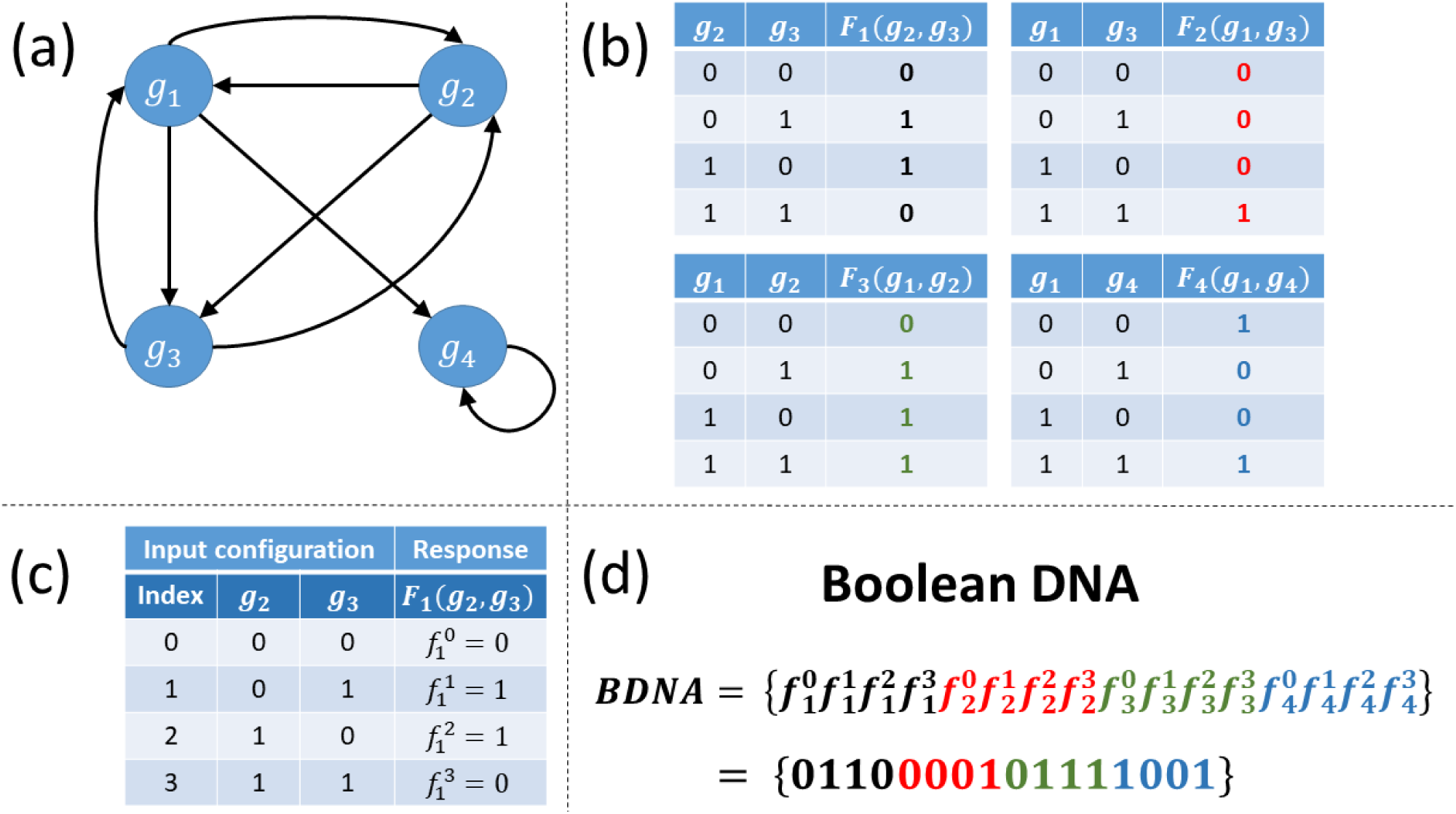
**(a)** Illustrative example of a network with 4 genes, each one having *K* = 2 inputs. The arrows indicate the regulatory interactions. **(b)** Specific example of the Boolean functions associated with each of the four genes in the network. **(c)** Structure of a Boolean function. The first column represents the index of the input configuration, ranging from *i* = 1 to *i* = 4. The second and third columns show the configurations of the inputs. The last column is the value of the function acquired on the *i*^*th*^ configuration of the inputs. **(d)** The Boolean DNA is constructed by concatenating the values of all the Boolean functions over all of their configurations. The genotype *G* of the network is the particular sequence the Boolean DNA consists of.

Once the inputs and Boolean functions have been assigned to every gene in the network, they do not change throughout time. What changes, is the state of expression of each gene according to Eq.(1). In other words, the inputs to each gene and its Boolean function are randomly assigned at the beginning of the simulation, when the network is constructed. After that, the topological structure of the network (the specific regulators of each gene) and the Boolean functions do not change in time. This is important since, as the network structure does not change throughout time, we can define a “Boolean DNA sequence” (BDNA) that characterizes the network. For that, let us note that a Boolean function with *K* inputs (arguments) has 2^*K*^ values, one for each configuration of its inputs (Fig. 1b). These 2^*K*^ configurations can be indexed according to their Boolean value, as Fig.1c shows. Thus, for instance, a function with *K* = 3 inputs has 2^3^ = 8 configurations, ranging from configuration {000} (the three inputs are off) to configuration {111} (the three inputs are on), with all the intermediate configurations in between. As Fig.1c shows, these configurations can be indexed from *i* = 1 to *i* = 2^*K*^. Let us define 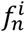 as the value of the *n*^*th*^ Boolean function *F*_*n*_ on the *i*^*th*^ configuration of its inputs (see Fig.1c). Then, the Boolean DNA of the network is defined as the concatenation of the values of all the Boolean functions in the network for all of their input configurations (see Fig. 1d):

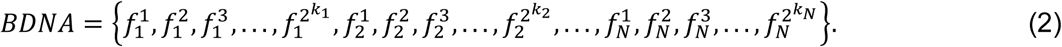

A network with *N* genes, with the *n*^*th*^ gene having *k*_*n*_ inputs, has a Boolean DNA with 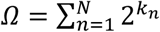 digits or loci. If *k*_*n*_ is the same for all the genes, let’s say *k*_*n*_ = *K* for every *n* = 1,2, …, *N*, then *Ω* = *N* × 2 ^*K*^. The dynamics of the network are determined by both its structural topology and its Boolean DNA. However, given a particular network structure, the Boolean DNA completely determines the dynamics of the network. Mutations in the Boolean DNA can change the network dynamics and its gene expression pattern. From now on we will indistinctly refer to the Boolean DNA of the network defined in Eq. 2 also as the *genom*e of the network, and to a particular Boolean sequence as its *genotype* (see Fig. 1d).

## 3 Genetic variants in a population

Since we want to study the heritability of phenotypes in a population, we have to define the phenotype in a quantitative way as well as the variants of that phenotype. To do this, let us consider a network with *N* genes connected in a specific way and with a specific Boolean DNA (see Fig.2). This network, which we will denote as *H*_0_, represents the genome (or part of the genome) of a given organism. Under some environmental conditions a subset of *N*_*s*_genes of this network, which we will refer to as the *transduction nodes* and represent by 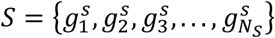, have an expression pattern determined by the function *R*_0_(*t*), defined over a time interval 0 ≤ *t* ≤ *T*. A gene expression pattern is always associated with a phenotypic trait. Therefore, we will refer to *R*_0_(*t*) as the *phenotype* of the network. This function may characterize the response of the transduction genes 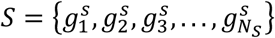 to external metabolites, stressors or environmental conditions (see Fig. 2). It can be for instance the phenotype needed to metabolize sugar or fatty acids. Now imagine that the environment permanently changes, and for the organism to adapt to the new environmental challenge, the subset of genes *s* has to change its expression pattern to a new function *F*(*t*). This new function *F*(*t*) is the *desired phenotype* for the organism to properly survive in the new environment. Therefore, the *actual* phenotype, encoded in the gene expression pattern *R*_0_(*t*), has to transform into the desired phenotype *F*(*t*). This change will not happen immediately after the environmental change has occurred. Instead, it will be through a series of mutations and partial adaptations that the actual phenotype *R*_0_(*t*)will transform into the desired phenotype *F*(*t*). Therefore, we will have to train the network *H*_0_ to perform the desired function *F*(*t*). To do this, we will implement a standard evolutionary algorithm in which a population of networks evolves through mutations and selection so that at each generation, only the networks that better approach the target function *F*(*t*) are the ones that survive and pass to the next generations.

**Fig. 2.**
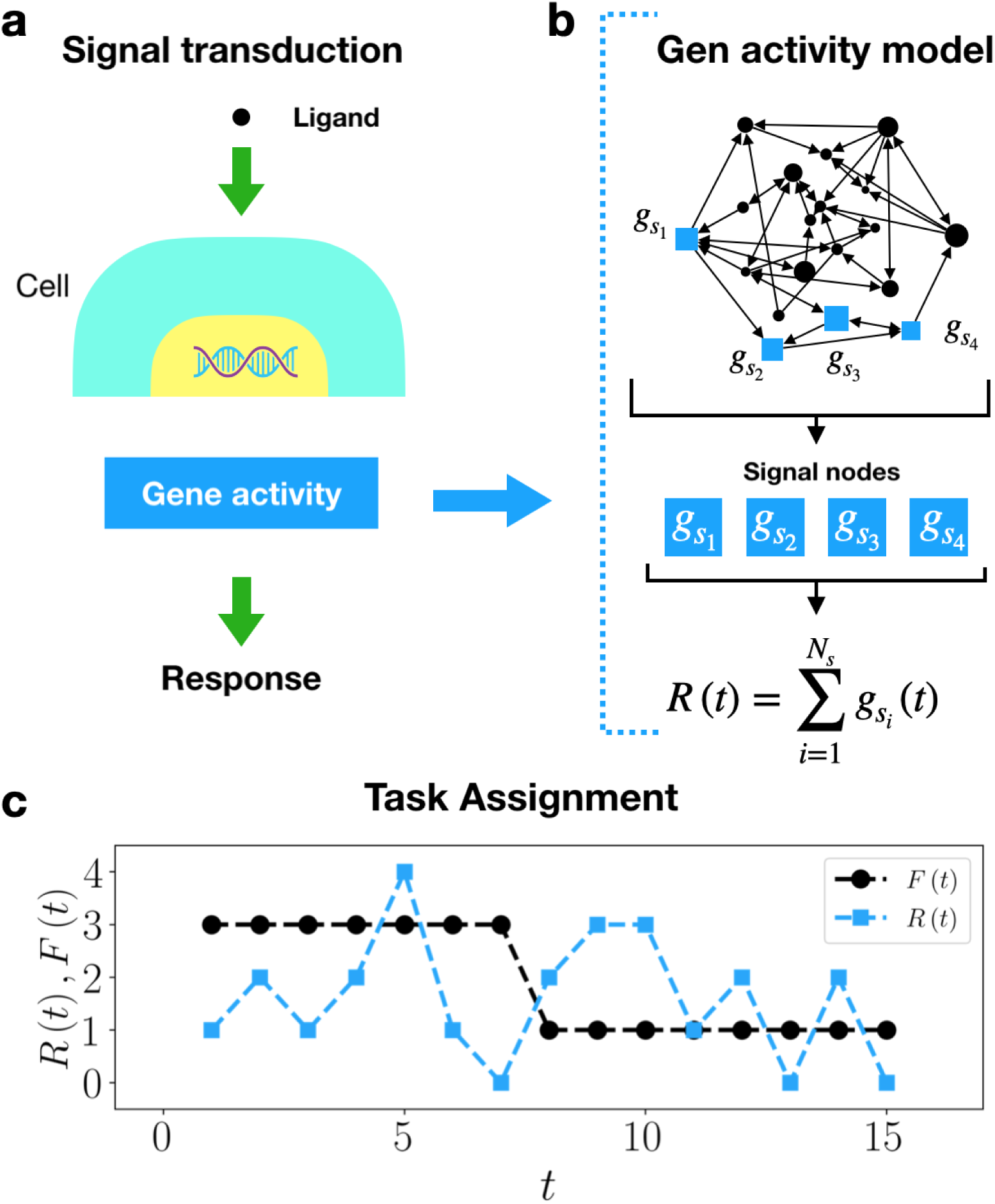
**(a)** Under external signals, a subset of the genes in a cell has to change its expression pattern in order to adapt to the environmental conditions. **(b)** A network representing the genome (or part of the genome) of a cell. The square nodes (in blue) are the transduction nodes that will respond to the external signal (a metabolite, stressor, etc.). These nodes generate a pattern of expression determined by the function *R*(*t*), which in this work is just the simple sum of the state of the nodes. **(c)** The desired phenotype *F*(*t*) is an arbitrary function defined over an interval of 0 < *t* ≤ *T*. In general, for a randomly constructed network, its actual phenotype *R*(*t*) is quite different from the desired phenotype *F*(*t*).

The function *R*_0_(*t*) is the phenotype of the network *H*_0_, and is determined by both its structure and its Boolean DNA (i.e., its genotype). The phenotype *R*_0_(*t*) has to transform into the desired phenotype *F*(*t*). To implement this transformation we will produce mutations in the Boolean DNA by keeping the structural topology of the network fixed. We measure the adaptation of the network to the new environmental conditions through the mean squared error *ξ*_0_ between the actual phenotype *R*_0_(*t*) and the desired phenotype *F*(*t*):

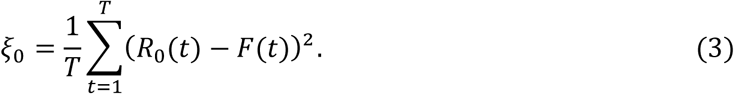

If *ξ*_0_ = 0, the network is perfectly adapted to the new task, as the actual phenotype *R*_0_(*t*) of the network is identical to the desired one *F*(*t*). By contrast, large values of *ξ*_0_ correspond to a poor adaptation, and the larger the value of *ξ*_0_, the poorer the adaptation.

Since the network *H*_0_ was constructed randomly, its phenotype *R*_0_(*t*) will, in general, be quite different from the desired phenotype *F*(*t*) (see Fig. 2c). For *R*_0_(*t*)to approach *F*(*t*) we have to mutate the genome of the network. This allows us to define the *variants* of the network as follows. Let us define *G*_0_ as the genotype of the network *H*_0_, namely, the particular Boolean sequence that characterizes the genome of *H*_0_ (like the particular Boolean sequence in Fig.1d). The variant *v*_*i*_ is a single mutation of *G*_0_ occurring at its *i*^*th*^ digit (locus). The variant *v*_*i*_ is thus a Single Nucleotide Polymorphism (SNP) occurring at the *i*^*th*^ locus of the genome. We will denote as *G*(*v*_*i*_) the genotype that contains the variant *v*_*i*_. In other words, *G*(*v*_*i*_) is a Boolean sequence almost identical to the genotype *G*_0_ of the original network *H*_0_ except that it has a SNP at the *i*^*th*^ locus of the genome (see Fig. 3). Clearly, for a network with *N* genes, each having *k*_*n*_ input connections, there are 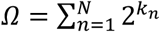 of such variants, *v*_1_, *v*_2_,…,*v*_*Ω*_ (see Fig.3). In this section and the following one, we will work with networks with *N* = 50 nodes, each with *K* = 2 regulators, which produces *Ω* = 200 loci in the Boolean DNA and the same number of variants.

**Fig. 3.**
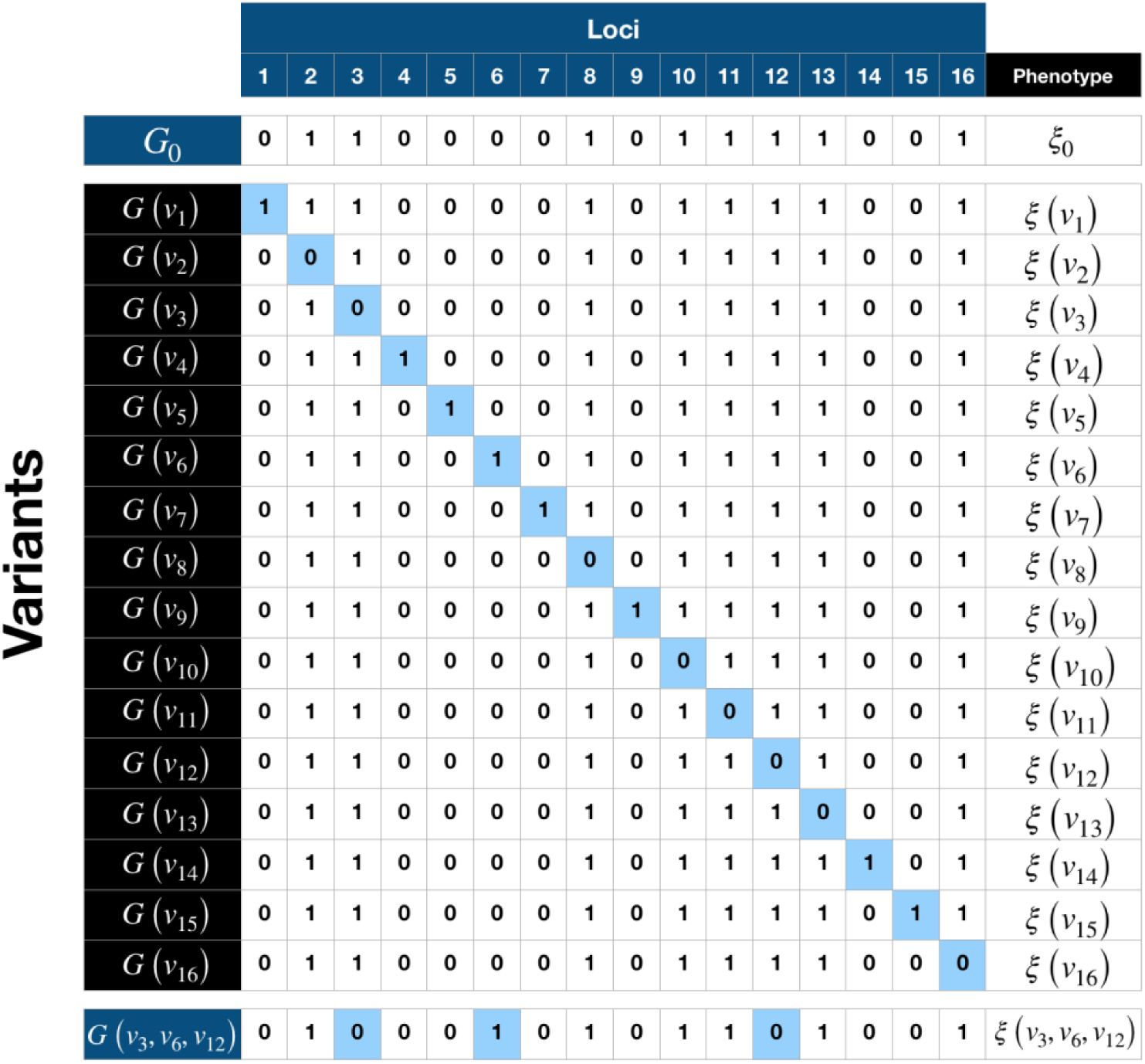
Illustrative example of the genetic variants for the genome of the network shown in Fig.1.This genome has *Ω* = 16 loci, indicated at the top of the matrix. The next row shows the genotype *G*_0_ of the original network *H*_0_ with respect to which the variants are defined. The subsequent rows show the particular genotypes *G*(*v*_*i*_) containing the variants *v*_*i*_, which are indicated in blue. Variant *v*_*i*_ consists of a single mutation (SNP) occurring at the *i*^*th*^ position (locus) of the genome *G*_0_ of the original network *H*_0_. The very last row shows a genotype *G*(*v*_3_, *v*_6_, *v*_12_) that contains the three variants *v*_3_, *v*_6_ and *v*_12_. The column to the far right just illustrates that each variant *v*_*i*_ corresponds to a particular error (phenotype) *ξ*_*i*_.

We have pointed out before that changes in the Boolean DNA of the network may change its dynamics. Therefore, even one mutation in the genotype of the network can change its phenotype *R*_0_(*t*). Therefore, to each variant *v*_*i*_ there corresponds a particular phenotype *R*_*i*_(*t*). Let us denote as *ξ*(*v*_*i*_) the error associated with the variant *v*_*i*_, which is defined in an analogous way as in Eq. 3:

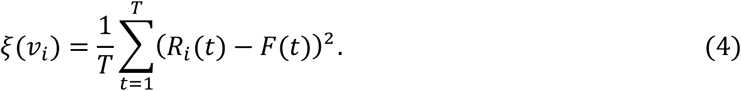

It should be noted that *ξ*(*v*_*i*_) is a quantitative measure of how well the phenotype *R*_*i*_(*t*) approximates the desired phenotype *F*(*t*). For instance, if *F*(*t*) were the phenotype required to metabolize sugar, then *ξ*(*v*_*i*_) ≈ 0 would mean that the organism with the genotype *G*(*v*_*i*_) is healthy, whereas a large value of *ξ*(*v*_*i*_) would mean that variant *v*_*i*_ is associated with diabetes. Different variants may produce different values of the error *ξ*(*v*_*i*_), which could be interpreted as variability in the diabetes phenotype. Therefore, we can consider *ξ*(*v*_*i*_) as the quantitative measure of the phenotype *R*_*i*_(*t*) that corresponds to the genotype *G*(*v*_*i*_), which ultimately is associated with variant *v*_*i*_. Since the error function *ξ*(*v*) is a quantitative measure of the phenotype, (ot how well the network performs the desired phenotype), we will indistinctly refer to *R*(*t*) and *ξ*(*v*) as the phenotype of the network.

Analogously, we will represent as 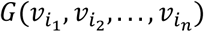 the genotype obtained from *G*_0_ by simultaneously implementing the *n* variants 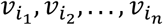, which are SNPs occurring at positions *i*_1_, *i*_2_, …, *i*_*n*_ of the genome (see Fig. 3). The phenotype corresponding to this genotype will be denoted as 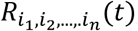, and its corresponding error as 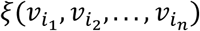, which is computed in a similar way as in Eq. 4.

Some of the variants *v*_1_, *v*_2_,…,*v*_*Ω*_ will have an associated error larger than *ξ*_0_ (the error of the original network *H*_0_), while some other variants will have an error smaller than *ξ*_0_. To measure the effect of the variant *v*_*i*_ on the adaptation of the network to the desired phenotype *F*(*t*), we define the error difference *δξ*_*i*_ as

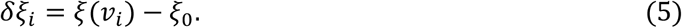

If *δξ*_*i*_ < 0 then the network containing the variant *v*_*i*_ is better adapted to the desired phenotype than the original network *H*_0_, whereas the opposite happens when *δξ*_*i*_ > 0. When *δξ*_*i*_ = 0 then variant *v*_*i*_ is neutral. Analogously, we can define the effect of the genotype 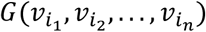 (with *n* variants) on the adaptation of the network as 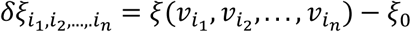. Fig.4a shows a plot of the effects *δξ*_*i*_ for all the variants *v*_1_, *v*_2_, …, *v*_200_ for a network with *N* = 50 and connectivity *K* = 2, while Fig. 4b shows the same data ordered in the increasing order of *δξ*_*i*_. Note that, while most of the variants are neutral, some variants produce considerably large effects (up to 50%), both positive and negative. This means that only one SNP can bring the network very close to the desired phenotype *F*(*t*), or very far from it.

**Fig. 4.**
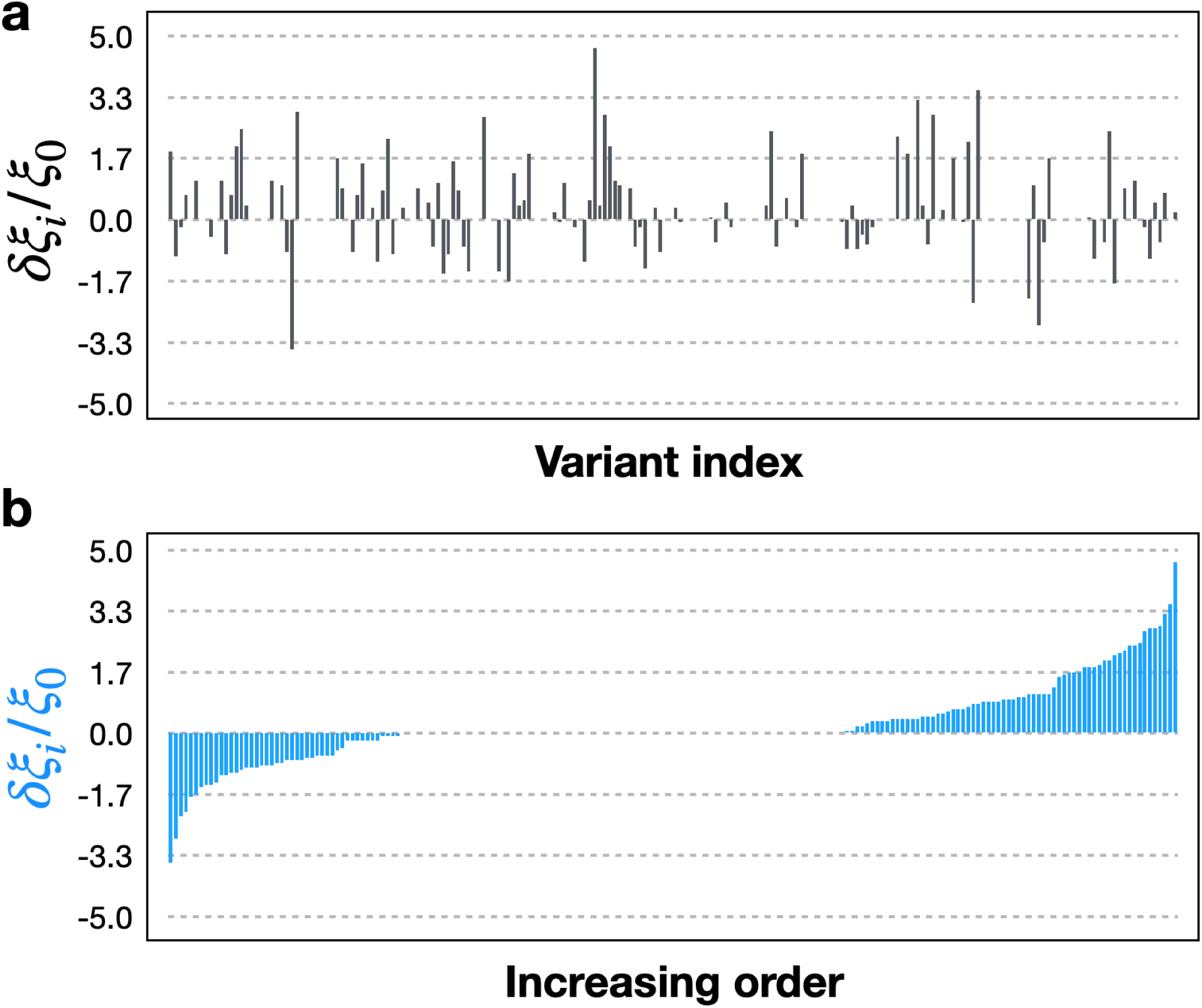
**(a)** Plot of the normalized effect *δξ*_*i*_/*ξ*_0_ of the variants *v*_*i*_ as a function of the variant index *i*, with 1 ≤ *i* ≤ 200, for a network with *N* = 50 and connectivity *K* = 2. **(b)** Same data as in the previous panel but ordered in increasing order. Note that some variants can have relatively large effects, ranging from -33% to almost 50% of the original error *ξ*_0_. Note also that most of the variants are neutral, with *δξ*_*i*_ ≈ 0.

One of the main arguments trying to explain the missing heritability of phenotypes is that the effects of the different variants on the phenotype are not additive [40, 11, 12]. Nonlinear interactions between these variants, known as epistatic effects, could be hiding the heritability of phenotypes. While this may be the case, here we show that nonlinear interactions between the variants are not enough to explain the missing heritability, for even when these epistatic interactions are negligible, the missing heritability persists.

## 4 Nonlinear (epistatic) effects

In the model proposed by Yang et al. in 2010 (Ref. [38]; see also the article by Eskin in Ref. [11]), to measure the heritability of a quantitative phenotype, linear regression is performed based on the following equation

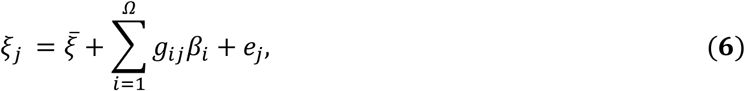

where *ξ*_*j*_ represents the value of the phenotype of the *j*^*th*^individual in the population, 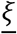 is the average value of the phenotype over the entire population, *β*_*i*_ is the size of the effect of the *i*^*th*^ variant *v*_*i*_ on the phenotype, and *g*_*ij*_ is a matrix that contains the variants of each individual, so its entries acquire the values 0, 1 or 2 depending on whether none of the alleles of the *j*^*th*^individual have the variant *v*_*i*_, only one allele has it, or both alleles have the variant. Finally, *e*_*j*_ is an error inherent in the measurement of the phenotype that is associated with the effect of the environment. It is clear that Eq. 6 assumes that the effects of the different variants are additive. In this section, we test whether or not the additivity hypothesis is true for the network model we are analyzing.

We construct a population of networks by making *N*_*P*_ = 10,000 identical copies of *H*_0_, which is a random network with *N* = 50 nodes, each one having *K* = 2 regulators. Therefore, the Boolean DNA has *Ω* = *N* × 2^*K*^ = 200 loci. Then, with probability *q*_*i*_ we mutate the *i*^*th*^locus of the genome of each network in the population (1 ≤ *i* ≤ *Ω*). Thus, a mutation in the first position of the genome (variant *v*_1_) will occur in *N*_*P*_ × *q*_1_ networks in the population, a mutation in the second position (variant *v*_2_) will occur in *N*_*P*_ × *q*_2_ networks, etc. The probabilities *q*_1_, *q*_2_, …, *q*_*Ω*_are chosen randomly in the interval 0 < *q*_*i*_ ≤ *δ*_*v*_, where *δ*_*v*_is the upper bound for the mutation probabilities. By increasing the threshold *δ*_*v*_ more mutations will be produced in the genomes of the networks. The reason for choosing different probabilities for the different loci in the Boolean DNA is to allow the different variants *v*_1_, *v*_2_, …, *v*_*Ω*_ to occur with different frequencies in the population. Note that the genome of one network in the population can accumulate several mutations, i.e., it can contain several variants 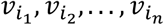. Figs. 5a and 5b show the probability *f*(*n*) that the genotype of a randomly chosen network in the population has *n* variants, for *δ*_*v*_ = 0.025 and *δ*_*v*_ = 0.1, respectively. In the first case the average number of variants in each genome is 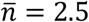 whereas in the second case it is 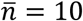.

**Fig. 5.**
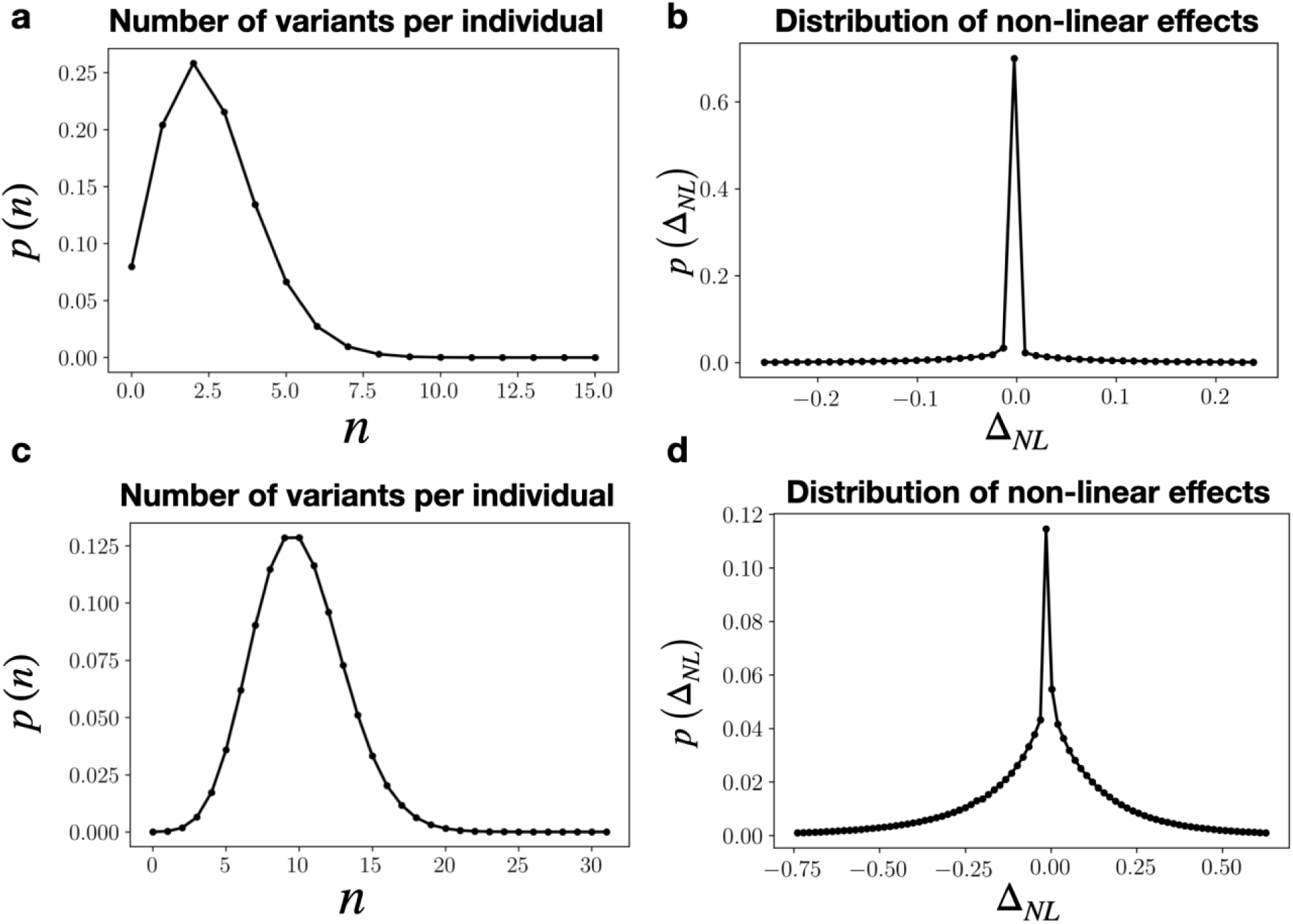
Distribution *P*(*n*) of the number of variants *n* per genome in the population for a variant probability threshold *δ*_*v*_ = 0.025 (panel **a**) and *δ*_*v*_ = 0.1 (panel **c**). In the first case, the average number of variants per individual is 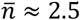, whereas in the second case it is 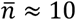. Panels **b** and **d** show the distribution *P*(*Δ*_*NL*_) of nonlinear effects *Δ*_*NL*_ for the corresponding populations in the previous two panels. Note that in both cases *P*(*Δ*_*NL*_) has a sharp maximum at *Δ*_*NL*_ = 0, which shows that in our model, the epistatic (nonlinear) interactions between individual variants are not important.

To determine the nonlinear interaction of the variants we consider a genotype 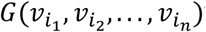 that has *n* variants. We know the error 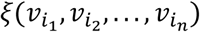 associated with this genotype because we can measure it. Analogously, we know the individual errors 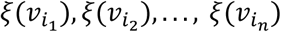 associated with each of the individual variants 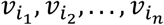. To determine whether these individual variants contribute linearly or not to the phenotype 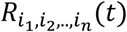 that corresponds to the genotype 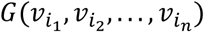, we define the *linear contribution* 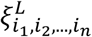 of the individual variants 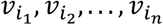 as

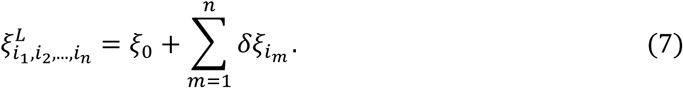

The linear contribution 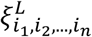 is an approximation to the real phenotype 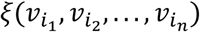 produced by the genotype 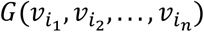. If the nonlinear interactions between the individual variants 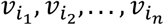 are indeed very strong, then 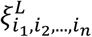 will be very different from 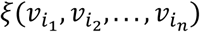, whereas if these nonlinear interactions are not so strong, then 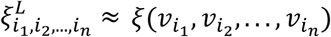. Therefore, a measure of the nonlinearity of the interactions between different variants is determined by the quantity

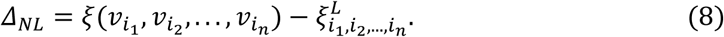

Figs. 5b and 5d show the probability distribution function *P*(*Δ*_*NL*_) for the same populations used to generate the data in Figs. 5a and 5c, respectively. Note that in both cases *P*(*Δ*_*NL*_) has a sharp maximum centered at *Δ*_*NL*_ = 0,which means that the nonlinear interactions between individual variants are not quite important in our model. This is particularly true for the case in which there is just a small number of SNPs per genome, as in Fig.5a, with an average of 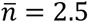 variants in each Boolean DNA. This represents a SNP occurrence probability of 0.0125 per locus (2.5/200). In the human genome this probability is even lower by at least one order of magnitude, as the SNP occurrence probability in the human genome is 0.0007 per nucleotide (on average 7 SNPs per 10 Kb [1, 27]). We do not claim that the results reported in Fig.5 can be extrapolated to the human genome (or to any other organism). In our model, and just in our model, the effect of most of the variants can be very well reproduced by a linear contribution of the effects of the corresponding individual variants. This is a remarkable result given that the network dynamics determined by Eq. 1 are highly nonlinear.

## 5 GWAS

For the sake of clarity, it is useful to keep in mind the analogy of the desired phenotype *F*(*t*) as the capability to metabolize sugars. Taking into account that the error function 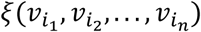 is a quantitative measure of how well the organism with the genotype 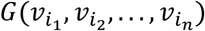 expresses the desired phenotype, then *ξ* = 0 would correspond to healthy organisms whereas *ξ* ≠ 0 would correspond to diabetic ones. In this analogy, we would be interested in the variants associated with this particular disease, namely, the variants that most increase the value of the phenotype *ξ*. (From Fig.4 it is clear that there is a symmetric behaviour between the positive and negative values of *δξ*; therefore, equivalent results would be obtained if we were looking for the variants that most decrease the value of *ξ*). In our model, we can exactly measure the value of the phenotype *ξ*(*v*_*i*_) corresponding to each variant *v*_*i*_, as well as the phenotype 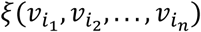 associated to any combination of variants 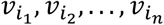. However, in real life we cannot know the phenotype *ξ*(*v*_*i*_) that each individual variant *v*_*i*_ produces because such variants generally do not occur isolated in the genome. Therefore, one has to estimate the contribution *ξ*(*v*_*i*_) that each individual variant *v*_*i*_ has on the phenotype under analysis. This allows us to identify the variants that are relevant to the phenotype in question. Those are variants that appear with a significantly large frequency among the population that presents the phenotype. This is where the Genome Wide Association Studies (GWAS) enters into play. The main objective of GWAS is to determine the variants that are significantly associated with the phenotype of interest. The main result of GWAS is to associate a *p*-value *p*(*v*_*i*_) to each variant *v*_*i*_. The variants with the smallest *p*-values will be the ones that are significantly associated with the phenotype. The natural question arises: how small the *p*-value has to be for the corresponding variant to be associated with the phenotype? As a rule of thumb, people in the community always choose a threshold *p*_*T*_ = 0.05for the *p*-value to be significant. If *p*(*v*_*i*_) < *p*_*T*_, then the corresponding variant *v*_*i*_ is accepted. Otherwise, it is rejected. Another fact to keep in mind is that, in practice, to perform a GWAS analysis not every individual in the population is tested. It is only a small fraction of the population whose DNA variants are analyzed. The size of the sampled population may have an important influence in the GWAS results, a phenomenon known as the *sampling size effect* [12, 11]. Some authors claim that by increasing the sampling size (the number of tested individuals), the results GWAS yields would be more trusty.

To simulate a GWAS analysis in our model, we consider a population consisting of *N*_*P*_ = 10,000 networks, each with *N* = 50 and connectivity *K* = 2. Variant *v*_*i*_ has been implemented with probability *q*_*i*_ in the genotype of the networks, with 0 < *q*_*i*_ < *δ*_*V*_ = 0.025 and 1 ≤ *i* ≤ 200. This would result in a population similar to the one used to generate the data reported in Figs. 5a and 5c. To perform GWAS on this population, we extract a subpopulation with *N*_*sub*_ = 500 networks, which are the ones that have the largest phenotypes *ξ* (in our analogy, these networks would correspond to clear cases of diabetes). In this subpopulation, the frequency 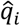 corresponding to the variant *v*_*i*_ may be different from the frequency *q*_*i*_ of this same variant in the original population. If 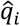 is significantly larger than *q*_*i*_, then one can think that *v*_*i*_ is a variant associated with the phenotype. By contrast, if 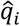 is significantly smaller than *q*_*i*_, then one can conclude that *v*_*i*_ is not associated with the phenotype. GWAS provides, through the *p*-value, the level of significance of the over-representation (or under-representation), of the variant *v*_*i*_ in the subpopulation that exhibits the phenotype.

Fig. 6a shows the normalized phenotype *δξ*_*i*_/*ξ*_0_ = (*ξ*(*v*_*i*_) − *ξ*_0_)/*ξ*_0_ in increasing order for a subpopulation *N*_*sub*_ = 500 networks and *Ω* = 200 variants, while Fig.6b shows the *p*-values corresponding to these variants. Clearly, the GWAS analysis in our model is detecting the variants *v*_*i*_ with the highest values of the phenotype *ξ*(*v*_*i*_) (the larger the value of *ξ*(*v*_*i*_), the smaller the corresponding *p*-value).

**Fig. 6.**
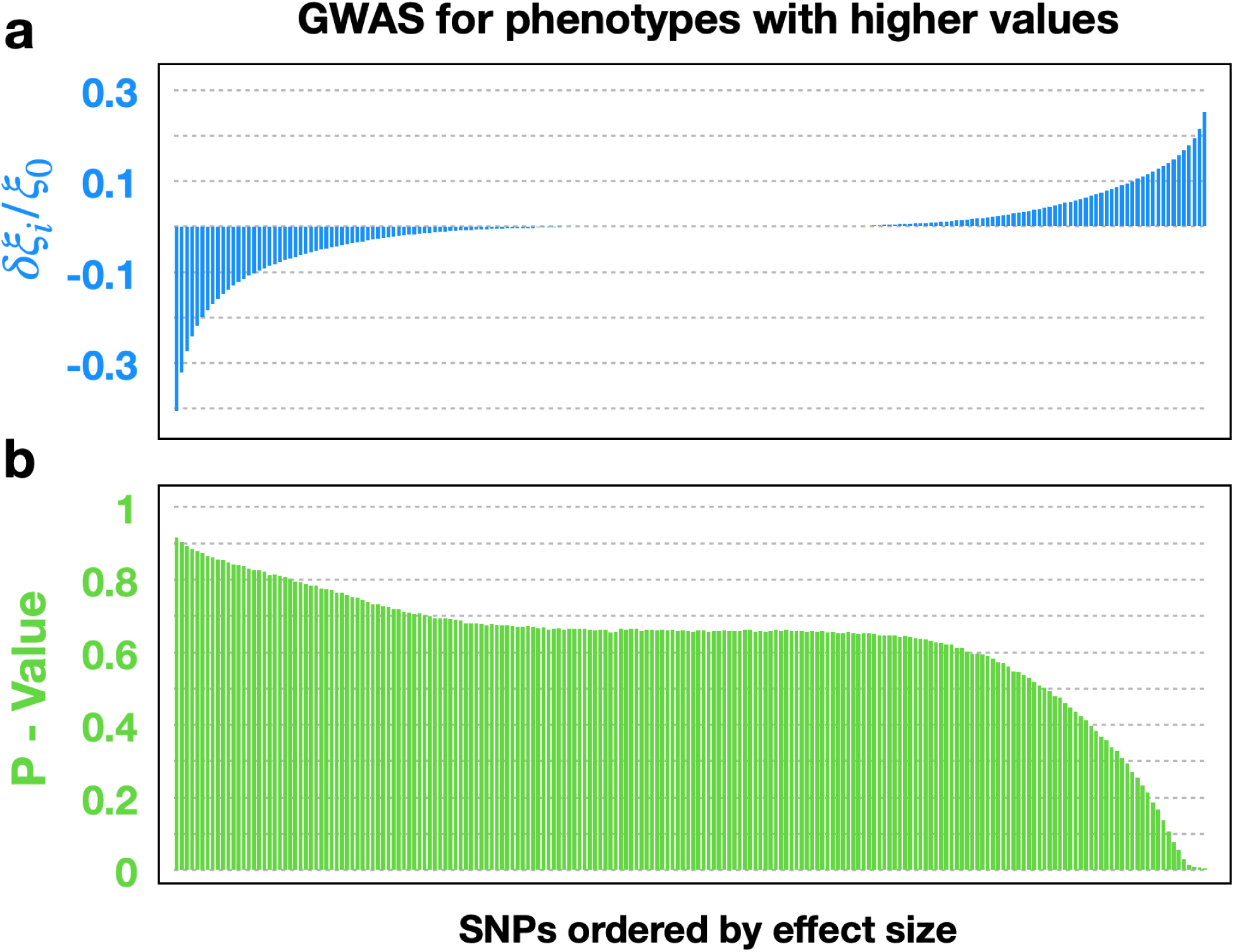
**(a)** Effect size *δξ*_*i*_ for the 200 variants *v*_*i*_ in a population of *N*_*P*_ = 10,000 networks, each with *N* = 50 nodes and connectivity *K* = 2. The data are plotted in the increasing order of *δξ*_*i*_. **(b)** A subpopulation of *N*_*sub*_ = 500 neworks (5%) is extracted. These networks are the ones with the largest effect sizes *δξ*_*i*_. The graph shows the *p*-value, computed through a GWAS analysis, that correlates the occurrence frequency 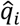 of variant *v*_*i*_ in the subpopulation with the corresponding effect size *δξ*_*i*_. It can be seen that the lowest *p*-values correspond to the largest effect sizes, which indicates that GWAS is correctly detecting the variants that contributed the most to the phenotype.

The results presented so far indicate two important aspects of our model: (1) epistatic effects (nonlinear interactions) are negligible when the number of variants in the genome is small, and (2) GWAS effectively detects the variants significantly associated with the phenotype under consideration (in this case, large values of the error *ξ*(*v*_*i*_)). In the next section we will show that the missing heritability problem is not necessarily a consequence of nonlinear effects nor of undersampling in the GWAS analysis. Instead, it may be a consequence of not taking into account the microbiota of the organism in the computation of the heritability, as the microbiota can be fundamental for the occurrence of some phenotypes.

## 6 Holobiont evolution and the missing heritability problem

So far we have not implemented any evolutionary algorithm to train the network *H*_0_to perform the desired task *F*(*t*). We have just generated genetic and phenotypic variability in a population of networks (initially all of them identical to *H*_0_) by implementing by hand the different variants *v*_1_, *v*_2_,…, *v*_*Ω*_ with some given probabilities. In this section, in order to compute the heritability of the phenotype, we implement an evolutionary algorithm to actually train (evolve) the network *H*_0_ to perform the desired task *F*(*t*). In what follows we will refer to *H*_0_ as the *host networ*k, in analogy with the host organism in a holobiont.

Following Huitzil et al. [16], the training of the host network *H*_0_ will be assisted by another network, *M*, which we will call the *microbial networ*k. This means that during the evolutionary process, the host network can acquire regulatory connections from the microbial network and vice versa (see Fig. 7). The main difference between the host and microbial networks are their mutation rates, with a mutation rate *μ*_*h*_ = 0.001 for the host network and *μ*_*m*_ = 0.01for the microbial network. The reason for this is that microbes can generate variability at a rate that is at least ten times larger than for the cells in a host organism (plants or animals). In our model, the host network *H*_0_ represents the host organism, whereas the microbial network *M* represents its microbiota. The regulatory connections between *H*_0_ and *M* represent the metabolic and genetic interactions between the host and its microbiota. Therefore, we will refer to the entire network made up of *H*_0_ and *M* as the *holobiont* [39]. It is important to mention that, although the host network is interacting with the microbial network, the target function *F*(*t*) is defined only for the host network and therefore it is just the phenotype *R*_0_(*t*) of the host network that will be used to train it. Therefore, the microbial network can be considered only as an auxiliary mechanism to help the host network reach its goal. The details of the evolutionary algorithm can be found in [16]. The important thing to mention here is that we have shown that the training of the host network is faster and more efficient with the help of the microbial network than without it and that most of the variability of the holobiont resides in the microbial network (due in part to its increased mutation rate).

**Fig. 7.**
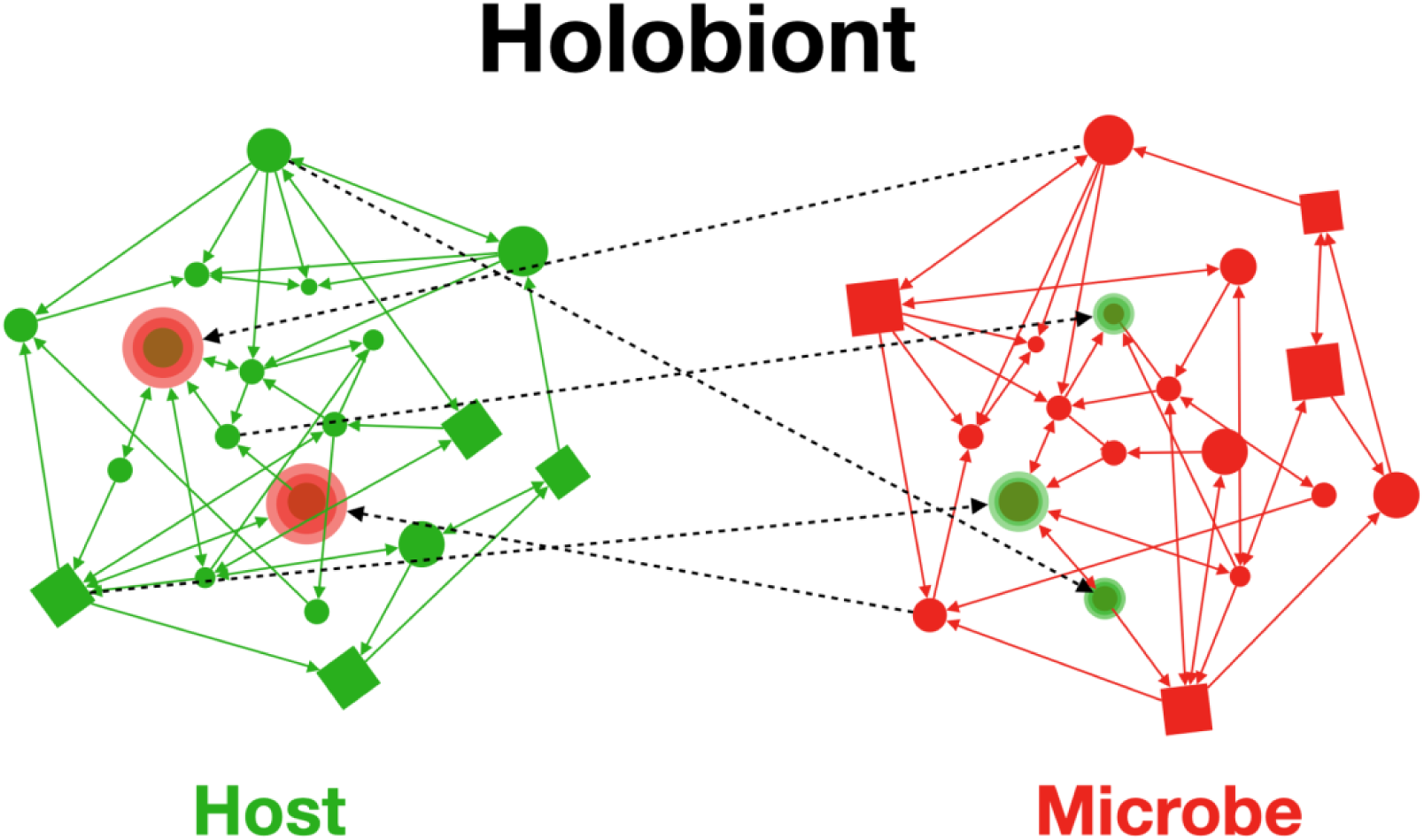
Schematic representation of a holobiont in our model. The holobiont consists of a host network and a microbial network, which can interact through regulatory connections between them (broken lines). In this example, the host network receives two regulatory connections from the microbial network, whereas the microbial network receives three from the host network. The host network still has to evolve to acquire the desired phenotype *F*(*t*), as in Fig. 2. The difference now is that the microbial network will “help” in the evolution of the host network.

The main idea in this section is to compute the heritability of the phenotype *R*(*t*) (which is quantitatively measured through the error function *ξ*) in two different ways: (i) by computing the genetic variability of the host network only, and (ii) by computing the genetic variability of the whole holobiont (host and microbial networks). As we will see, the missing heritability is considerably larger in the first case than in the second one.

The evolutionary model consists of a population of *N*_*P*_ = 1000 holobionts. Initially, the host and microbial networks in each holobiont have *N* = 50 nodes, each node with *K* = 2 regulatory connections. At each generation, each node in each network is mutated with probability *μ*_*h*_ for the host network and *μ*_*m*_ for the microbial network. Once a node has been chosen for mutation, the mutations consist of the following: (i) Adding a new regulatory interaction (input connection); (ii) rewiring an existing regulatory interaction; (iii) changing the value of one entry of the Boolean function. Mutations (i) and (ii) can occur between nodes within the same network, or between one node in the host network and the other node in the microbial network. This last possibility is what makes the two networks develop regulatory interactions between them (see Fig. 7).

Let us denote as *R*_*n*_(*t*) and *ξ*_*n*_ the phenotype of the *n* ^*th*^ holobiont in the population and its corresponding error, respectively. At each generation, we choose the best 100 holobionts in the population (the ones with the lowest values of *ξ*_*n*_) to pass to the next generation. Then, we replicate these holobionts by making 10 copies of each one in order to restore the population to its original size *N*_*P*_ = 1000. Then we repeat the entire process (mutation, selection, replication, etc.). We do this for several generations until the average population error 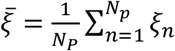 becomes smaller than a threshold *ε* = 1. This means that on average most of the holobionts in the population are well adapted to the desired phenotype *F*(*t*). We stop the simulation as soon as the condition 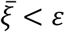 is satisfied. However, this algorithm eventually produces a population with almost no variability, namely, a population in which almost all the holobionts are identical, a phenomenon known as *purifying selection* [8, 20, 15]. Therefore, in order to generate variability in the population, we proceed as follows. After the condition 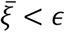 has been fulfilled, we continue the evolutionary process for 5 more generations. But now, in each one of these 5 generations we mutate only the Boolean DNA of the networks, which makes their corresponding errors to change: some errors will increase while some others will decrease. Then, at each one of these 5 generation, we allow the *n*^*th*^ holobiont to pass to the next generation with a probability *p*_*n*_ = *C*/*ξ*_*n*_ (*C* is a proportionality constant). Thus, holobionts with a large error (poorly adapted) still have a probability, although small, to continue through the next generation. In this way, the final population will have more variability than if we were always to choose the best holobionts and replicate them.

At the end of this process all the holobionts in the population have the same structural topology but different genotypes, all of the same length. (The genome of a holobiont is the concatenation of the genome of its host network and the genome of its microbial network). Since through the evolutionary process regulatory connections within and between the host and microbial networks were added or rewired, different genes in the final networks will have a different number of input connections. We will denote as *Ω*_*h*_, *Ω*_*m*_and *Ω*_*HL*_the length of the genomes of the host network, the microbial network, and the entire holobiont, respectively, with *Ω*_*HL*_ = *Ω*_*h*_ + *Ω*_*m*_. Since all the holobionts in the final population have the same structural topology but different Boolean DNAs, it is possible to align their genotypes and determine a consensus sequence in the same way as one does with the genetic sequences of real organisms (see Fig. 8). Once the consensus sequence of the population has been determined, the variants are defined as mutations (SNPs) of the consensus sequence. There are *Ω*_*h*_ variants for the host network, *Ω*_*m*_ variants for the microbial network, and of course *Ω*_*HL*_ = *Ω*_*h*_ + *Ω*_*m*_ variants for the holobiont, which we will denote as 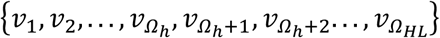. In the final population each variant *v*_*i*_ (*i* = 1, …, *Ω*_*HL*_) occurs with a frequency *q*_*i*_. Rare variants in the population are defined as those that satisfy the condition *q*_*i*_ < *δ*_*f*_, where *δ*_*f*_ is a parameter chosen in the interval [0.001,0.01]. We vary *δ*_*f*_ in this interval to analyze the effect of taking into account (or not) the less common variants in the computation of the heritability, finding no significant changes in the results when *δ*_*f*_varies in the interval mentioned above.

**Fig. 8.**
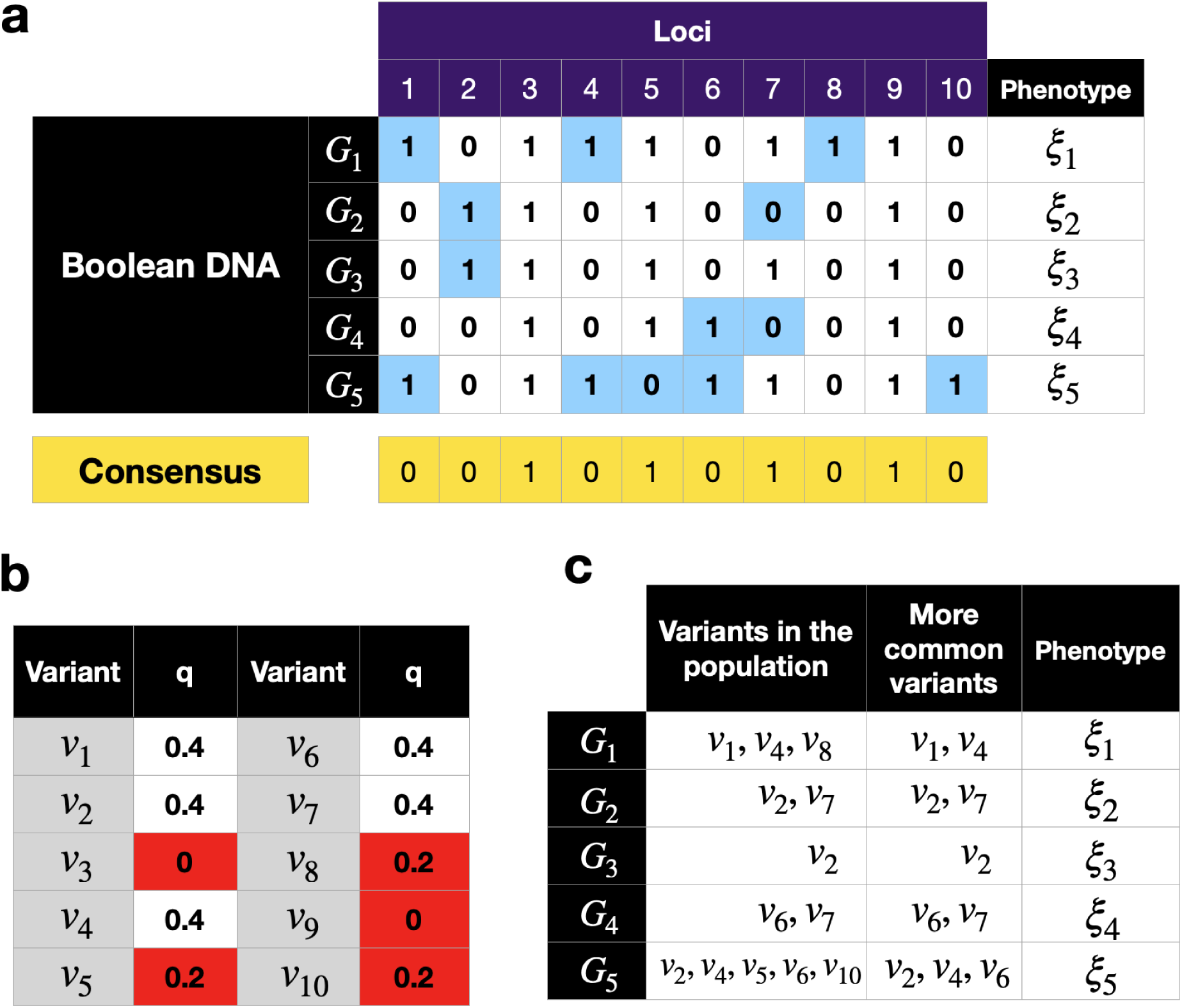
**a)** Illustrative example of five Boolean genotypes *G*_1_, *G*_2_, *G*_3_, *G*_4_ and *G*_5_ corresponding to five individuals in a population. Each genome has 10 loci. The consensus sequence, which is shown at the bottom of the table, is constructed using a simple majority rule at each locus. The variants (or SNPs) of each individual are those loci whose value is different from the corresponding one in the consensus sequence. These values are highlighted in blue in the figure. Since the genotypes of the 5 individuals are different from each other, their phenotypes *ξ*_*i*_ will usually be different from each other as well. **(b)** Table showing the occurrence frequency *q*_*i*_ of each variant *v*_*i*_ in the population. Less frequent variants are highlighted in red. **(c)** Knowing the consensus sequence of the population, all the information in (a) can be summarized by just indicating the variants of each individual.

The strict sense heritability *h*^2^of the phenotype is defined as [11,38]

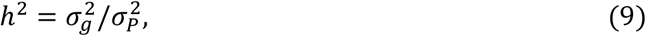

where 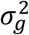 and 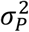 are the genotypic and phenotypic variances (or variabilities) of the population, respectively. The phenotypic variance 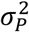 in our model is easily computed, as it is just the variance of the phenotype *ξ* over all the holobionts in the population:

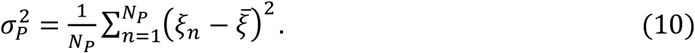

The genotypic variance 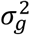 is more difficult to compute, as it has to do with the variants that are associated with the desired phenotype. Therefore, one has to determine first, though GWAS, the variants that correlate with the phenotype and then compute the variability of those variants throughout the population. Nonetheless, we can see that in the genetic variance 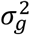 of our holobiont model there are two contributions, one coming from the genome of the host network and the other other from the genome of the microbial network. We will denote these contributions as 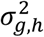 and 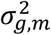, respectively. Therefore, 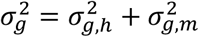 and the heritability in Eq.9 can be written as

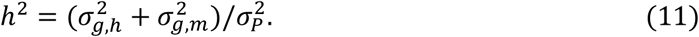

Defining the host heritability and the microbial heritability as 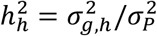 and 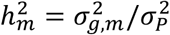, respectively, then the heritability of the holobiont is 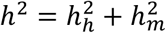. Since both 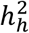 and 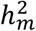 are positive, it is clear that the two following conditions are satisfied: 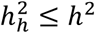 and 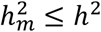. If when computing the genetic variance 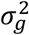 only the genome of the host organism is considered, as it is usually the case, then we will obtain a value 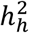 which could be considerably smaller than the real value *h*^2^, unless the microbial heritability 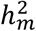 is negligibly small compared to the host heritability 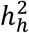. But this is not a sensible hypothesis, in view that the microbiota is strongly correlated with the emergence of some phenotypes (like diabetes). In our network model, the microbial network *M* strongly participates in the training of the host network *H*_0_ to achieve its desired phenotype *F*(*t*). The influence of the microbial network *M* on the training of the host network *H*_0_ is stronger the more regulatory connections occur between them. We will denote as *K*_*ex*_ the number of regulatory connections from the microbial network to the host network, and refer to them as the *external host regulations*.

The details of the algorithm to compute the heritability *h*^2^in our model is presented in the Appendix. It is based in Eq. 6 and closely follows the method reported in Eskin 2015 and Yang 2010 [38, 11]. The results of this computation are presented in Fig.9, which shows the host and microbial heritabilities, 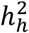 and 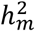 respectively, as functions of the number of external host regulations, *K*_*ex*_. These results were computed for the final population of holobionts obtained from the evolutionary algorithm described in this section. Note from Fig.9 that when *K*_*ex*_ = 0, i.e. when there is no interaction between the host and microbial networks, almost all the contribution to the heritability *h*^2^of the phenotype comes from the heritability 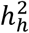 of the host network, as should be expected. However, when *K*_*ex*_ increases the interaction between the host and microbial networks becomes stronger. In this case, the heritability 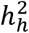 of the host network decreases while the heritability 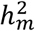 of the microbial network increases. Even for *K*_*ex*_ ≈ 2.5, which corresponds to a relatively weak interaction between the host and microbial networks, the microbial heritability 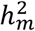 starts to surpass the host heritability 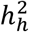. For *K*_*ex*_ = 10 (a strong interaction) almost all the contribution to the heritability *h*^2^ comes from the microbial network. In this strong-interaction case 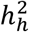 is very small with the result that 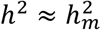.

**Fig. 9.**
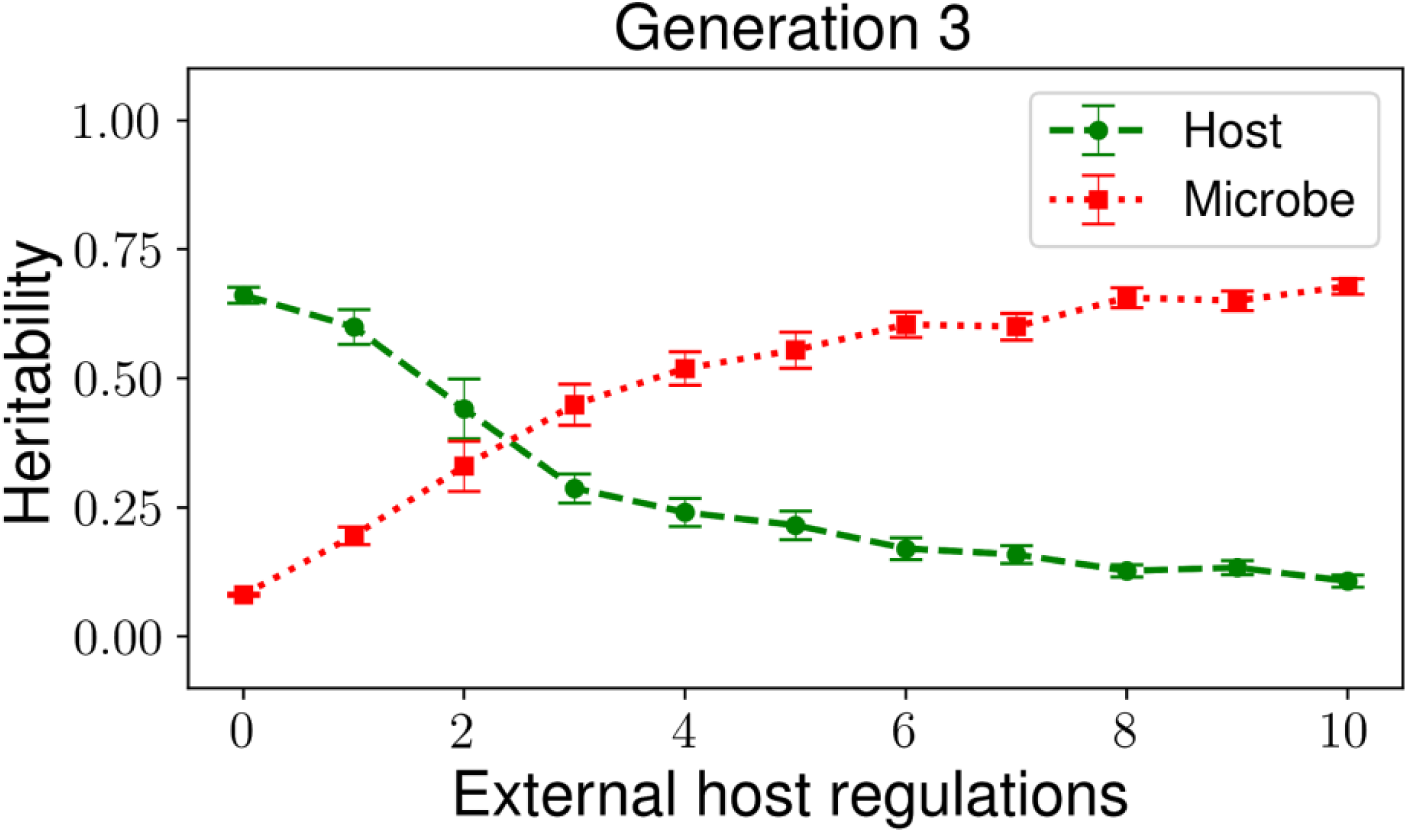
Host and microbial heritabilities, 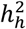 and 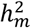 respectively, as functions of the number *K* of host external regulations. Only for small values of *K*_*ex*_ (weak host-microbe interaction) the heritability of the host network is larger than the heritability of the microbial network. However, as *K*_*ex*_ increases (strong interaction), 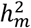 increases and very rapidly becomes larger than 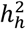.

## 7 Discussion and conclusions

The missing heritability problem has been a matter of intense debate for more than one decade. The fact that many phenotypes that are transmitted across generations cannot be significantly correlated with a particular set of genetic variants has generated many possible explanations. Among those explanations three sand out [12]. First, it has been proposed that epistatic (nonlinear) effects emerging from the interactions between these genetic variants may be hindering the identification of significant correlations between the genetic variants and the phenotype under consideration. Second, there is the undersampling problem, which consists in that rare variants that have a strong effect on the phenotype are not being detected due to the combination of two factors: these variants occur with very low frequency in the population and the size of the sampled subpopulation is too small. Third, there are many common variants whose combined effect on the phenotype is very strong, but whose individual effects are too small to be detected individually. While these answers to the missing heritability problem might be true, there is one aspect that has not been considered when measuring the heritability of the phenotype of a given organism, which is the genetic variability of its microbiota.

More than one decade ago, the pioneering work by Turnbaugh and his coworkers [31] showed that the microbiota can have a strong influence on the phenotypic traits of its host organism. This is particularly true for many of the phenotypes that researchers have tried to correlate with genetic variants, such as obesity, diabetes, cancer, metabolic syndrome, etc. It has also been shown that the microbiota can be transmitted across generations. Furthermore, there is evidence that the microbiota can evolve together with its host organism, strongly influencing (or even substituting) some of the genetic and metabolic functions of the host organism. Therefore, it is natural to assume that, when computing the heritability of some phenotypic trait, (particularly one that is strongly influenced by the microbiota), one does not only have to take into account the genotypic variance of the host organism, but also the genotypic variance of its microbiota. This is the approach that we have adopted in this work.

We have presented an evolutionary algorithm based on the Boolean gene regulatory network model proposed by S. Kauffman in 1969, which has proven to accurately reproduce the gene expression patterns experimentally observed for several organisms. The objective is to train a population of networks to perform a predefined task *F*(*t*), which represents the phenotype of the networks. To do this training, we mutate the Boolean functions of the networks (their genotypes), which introduces genetic variability in the population (all the networks have the same structural topology but slightly different Boolean functions). The main advantage of working with this model is that we can exactly measure the effect that each variant has on the phenotype, which in turn allows us to simulate a GWAS analysis in order to compute the correlations between the genetic variants and the desired phenotype. From this analysis, three important results are obtained. First, most of the variants have a very small effect on the phenotype, but there are a few variants that have a strong effect (Fig. 4). Second, when several variants are present in the genotype of one individual, their effects on the phenotype are mostly additive (Fig. 5). Therefore, epistatic (nonlinear) effects can be ignored when computing the contribution of several variants to the phenotype. This is an interesting result given that the dynamics determined by Eq. 1 are highly nonlinear. And third, GWAS effectively reveals the variants that have a strong effect on the phenotype (Fig. 6). These results indicate that all the conditions are met in our model to obtain a good estimate of the heritability *h*^2^ of the phenotype based on the variability of the Boolean DNA of the networks.

To compute this heritability we evolved a population of holobionts, where each holobiont consists of a host network and a microbial network that can interact through regulatory connections between them (Fig. 7). The goal of the evolutionary process is to train the host network to acquire the desired phenotype *F*(*t*). It is important to stress that the phenotype is defined only on the host network. However, since the host and microbial networks can interact, the adaptation of the host network to the desired phenotype also depends on the microbial network, and this dependence is greater the more regulatory interactions exist between the host and microbial networks. Therefore, in order to compute the heritability *h*^2^ of the phenotype, one has to take into account the genetic variance of both the host and the microbial networks. The heritability *h*^2^ has two contributions and can thus be written as 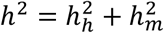, where 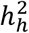 and 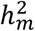 are the heritabilities computed by taking into account the genetic variance of only the host and only the microbial networks, respectively. It has happened that when the heritability of a particular phenotypic trait, (such as height, cancer or diabetes), is computed, only the genetic variance of the human genome is taken into account. Therefore, researchers all over the world have been computing 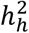 instead of *h*^2^. From Fig. 9 it can be seen that 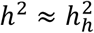 (i.e. 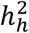 is a good estimate of *h*^2^) only when there are no interactions whatsoever between the host and the microbial networks (this would correspond to an organism with no microbiota). However, as soon as the number of interactions between these two networks starts to increase, very quickly one has 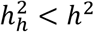. The difference 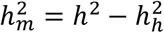 is the missing heritability, and is completely attributed to the genetic variability of the microbiota. As Fig. 9 shows, this missing heritability becomes larger the stronger the interaction between the host and the microbial networks.

The most important conclusion of the work presented here is the following: even when epistatic (nonlinear) effects between genetic variants can be neglected, and even when GWAS can efficiently detect the most important variants that contribute to a phenotype, if the genetic variance of the microbiota is not taken into account in the computation of the heritability *h*^2^ of some particular phenotype, then this heritability will always be underestimated by a large amount. This is particularly true for those phenotypes that are strongly determined by (or correlated with) the microbiota composition of the host organism. This problem has recently started to be addressed experimentally through Metagenome Wide Association Studies (MWAS), through which the genetic variance of both the host and its microbiota can be measured [35]. Our results strongly suggest that MWAS will be essential to fill the missing heritability gap.

## Acknowledgement

S.H. thanks CONACyT for a Ph.D. scholarship and the C3-UNAM for economic support. This work was partly supported by PAPIT-UNAM grant IN226917.

## Appendix

Following Yang 2010 and Eskin 2015 [38, 11], to measure the heritability *h*^2^ of a given phenotype, we start with Eq. 6 of the main text, which determines the value *ξ*_*j*_ of the phenotype of the *j*^*th*^ individual in the population as a linear contribution of the effects of the different genetic variants *v*_*i*_ (*i* = 1, 2, …, *Ω*_*HL*_):

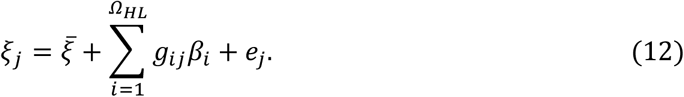

In this equation, 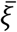 is the average value of the phenotype in the population; *β*_*i*_ is the contribution of variant *v*_*i*_ to the phenotype; *e*_*j*_is an error that has to do with the unknown effect of the environment on the phenotype; and *g*_*ij*_ is a matrix whose entries acquire the values 1 and 0 depending on whether the *j*^*th*^ individual contains variant *v*_*i*_ or not, respectively. For simplicity in the calculation, it is convenient to perform the change of variable

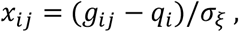

where *q*_*i*_ is the occurrence frequency of variant *v*_*i*_ in the population, and 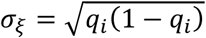 is the standard deviation of this quantity. With this change of variable, Eq.(12) can be written in matrix form as

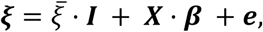

where 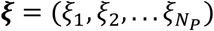 is the vector containing the phenotypes of the individuals in the population (analogously for the vectors ***β*** and *e*), and ***I*** is the identity matrix. In our model, we know exactly the effect *β*_*i*_ that variant *v*_*i*_has on the phenotype (it is the error difference *δξ*_*i*_ defined in Eq. 5 and reported in Fig.4). However, in a real situation this effect cannot be known accurately. Instead, it has to be estimated ad 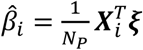, where ***X***_*i*_ is the *i*^*th*^ row of matrix ***X*** and *N*_*P*_ is the number of individuals in the population.

To determine whether the effect of variant *v*_*i*_ is or not correlated with the phenotype, we compute the *p*-value of the effect of this variant on the phenotype. The *p*-value *p*(*v*_*i*_) corresponding to the variant *v*_*i*_ is computed as

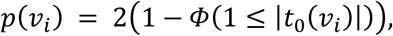

where 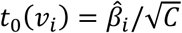. In this expression, 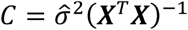 is the covariance of the linear regression with 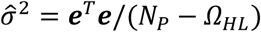 being the error of the estimation (*Ω*_*HL*_ is the number of different variants occurring in the population). The function *Φ*(*x*) is the (cumulative) normal distribution function.

Once the effect 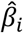 of each variant *v*_*i*_ has been estimated, the variants whose effect on the phenotype is significantly small are discarded. Let us denote as *s*_*V*_ the set of variants that are not discarded and remain in the analysis, namely, the set of variants that have a strong effect on the phenotype. These variants can occur in the genome of both the host network and microbial networks. Therefore, the set *s*_*V*_ can be partitioned into two disjoint subsets, *s*_*H*_ and *s*_*M*_ such that *s*_*V*_ = *s*_*H*_ ∪ *s*_*M*_, where *s*_*H*_ is the set of relevant variants that occur in the genome of the host network and *s*_*M*_ is the set of relevant variants occurring in the microbial network. The genetic variance of the genotype is then estimated as

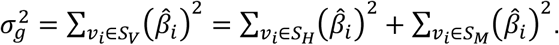

The heritability *h*^2^ is then computed as

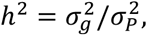

where 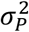 is the phenotypic variability computed as

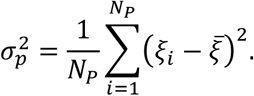

